# Finding divergent sequences of homomorphic sex chromosomes *via* diploidized nanopore-based assembly from a single male

**DOI:** 10.1101/2024.02.29.582759

**Authors:** Igor Filipović, John M Marshall, Gordana Rašić

## Abstract

Although homomorphic sex chromosomes can have non-recombining regions with elevated sequence divergence between its complements, such divergence signals can be difficult to detect bioinformatically. If found in genomes of e.g. insect pests, these sequences could be targeted by the engineered genetic sexing and control systems. Here, we report an approach that can leverage long-read nanopore sequencing of a single XY male to identify divergent regions of homomorphic sex chromosomes. Long-read data are used for *de novo* genome assembly that is diploidized in a way that maximizes sex-specific differences between its haploid complements. We show that the correct assembly phasing is supported by the mapping of nanopore reads from the male’s haploid Y-bearing sperm cells. The approach revealed a highly divergent region (HDR) near the centromere of the homomorphic sex chromosome of *Aedes aegypti*, the most important arboviral vector, for which there is a great interest in creating new genetic control tools. HDR is located ∼5Mb downstream of the known male-determining locus on chromosome 1 and is significantly enriched for ovary-biased genes. While recombination in HDR ceased relatively recently (∼1.4 MYA), HDR gametologs have divergent exons and introns of protein coding genes, and most lncRNA genes became X-specific. Megabases of previously invisible sex-linked sequences provide new putative targets for engineering the genetic systems to control this deadly mosquito. Broadly, our approach expands the toolbox for studying cryptic structure of sex chromosomes.

## Background

Homomorphic sex chromosomes are surprisingly common among animals and plants (Wright et al. 2016). Lack of cytological differences between the sex chromosome pair (homomorphy) can be due to their recent origin. Namely, after proto-sex chromosomes stop recombining around a newly-acquired sex-determining locus, the chromosome limited to the heterogametic sex (Y, W) tends to lose genes and accumulate repetitive and heterochromatic sequences over time (Charlesworth et al. 2005). However, many species have old homomorphic sex chromosomes, posing questions around mechanisms that prevent accumulation of substantial divergence between X and Y (or Z and W) (Stöck et al. 2011; Vicoso et al. 2013a, 2013b). Still, homomorphic sex chromosome pairs can contain divergent non-recombining regions extended beyond the sex-determining locus (Charlesworth 2017). The enhanced ability to accurately detect and characterize such regions is not only needed to refine our understanding of sex chromosome evolution, but has bioengineering applications. For instance, X- or Y-specific sequences are required for genetic pest management strategies that engineer genetic sexing systems and sex ratio distorters such as Y-linked X-shredders or X-poisoning gene drives (Gamez et al. 2021; Fasulo et al. 2020).

Regions of sex chromosomes that are either fully or partly differentiated are typically identified *via* next-generation short-read sequencing using the information on the depth of read coverage or allelic patterns from male and female samples (reviewed in (Palmer et al. 2019)). Short reads from both sex chromosomes (X and Y, or Z and W) that have low divergence are expected to map to the same regions of the haploid genome assembly. In that scenario, their sequence difference can be detected as the increased heterozygosity limited to the heterogametic sex (XY males, ZW females), or as different allele density and frequency between males and females (Fontaine et al. 2017; Pucholt et al. 2017). When divergent at many sites, Y-linked sequences become ‘invisible’ (null alleles) as they fail to align to the reference assembly. In that case, depth of read coverage across the divergent region is expected to be two times lower in the heterogametic sex compared to the homogametic sex or to the autosomal parts of the genome (Hall et al. 2013). However, this approach is sensitive to variations in methodology; the choice of a reference genome assembly or parameters for sequence alignment and filtering can make a difference between successfully recovering or completely missing the subtle signals of X-Y (or Z-W) divergence (Darolti et al. 2022).

Haplotype-resolved genome assemblies should enable precise characterization of divergent features of sex chromosome pairs, but they have been challenging to produce for homomorphic sex chromosomes. With low levels of nucleotide difference between X and Y (or Z and W), phasing their haplotypes with SNPs from short-read sequences is difficult (Browning and Browning 2011). Another challenge is often the repetitive and heterochromatic nature of a Y- (W-) linked sex-determining region, prohibiting its accurate assembly with short-read data (Tomaszkiewicz et al. 2017; Chen et al. 2012). Long-read sequencing can overcome these problems, but the higher error rate of platforms like Pacific Biosciences (PacBio) CLR and Oxford Nanopore Technologies (ONT) impedes the correct separation of X and Y (Z and W) haplotypes (Ebler et al. 2019). When parents and progeny are available for sequencing, an approach called trio binning can produce offspring’s haplotype-resolved long-read assembly with the help of highly-accurate short reads from two parental genomes (Koren et al. 2018). Such pedigrees, however, may be unavailable for many non-model organisms, making the single-sample assembly approaches preferable. Appearance of high-fidelity (HiFi) PacBio long reads, whose accuracy is comparable to the short-read technology (Hon et al. 2020), facilitated the assembly of fully or partially phased homomorphic sex chromosomes from a single specimen (e.g. XY eel (Xue et al. 2021), ZW strawberry (Cauret et al. 2022)). Here, we demonstrate an approach that can produce accurately phased divergent regions of homomorphic sex chromosomes with the help of ONT data from a single XY specimen.

We applied the approach to the male *Aedes aegypti* mosquito that was sequenced with the MinION device, a widely-accessible ONT platform. Following the initial construction and polishing, the assembly is diploidized in a way that maximizes sex-specific differences between its haploid complements. This process uses the statistic we call the Sex-Bias Score (SBS), which quantifies the differential mapping of short read data from males and females. Alignment of the protein sequences within the male-biased (Y-linked) contigs (SBS > 0) is used to identify X and Y gametologs in the diploid assembly. We also show that the correct X-Y phasing is supported by the mapping of the ONT reads from male’s haploid Y-bearing sperm cells.

*Aedes aegypti* mosquito is the most important global vector of arboviruses for which there is a great interest and effort to create new genetic control tools (Alphey et al. 2013). Its genome is large (1.3 Gb), riddled with repetitive elements and composed of three chromosomes (Matthews et al. 2018; Nene et al. 2007). Like all members of the Culicinae subfamily, it has an old homomorphic sex chromosome system, and its Y chromosome contains a male-determining locus (M-locus) with a male-determining “master switch”, the *Nix* gene (Hall et al. 2015). Y and X chromosomes are commonly designated as M and m chromosomes, respectively, with males having the M/m karyotype and females the m/m karyotype. In the current *Ae. aegypti* reference genome assembly (AaegL5.0, Matthews et al. (Matthews et al. 2018)), The M-locus is annotated as a small region (∼1.5 Mb-long) near the centromere of chromosome 1 (Matthews et al. 2018). Outside of the non-recombining M-locus, the sex-determining chromosome 1 is thought to behave in an autosomal-like manner and its sequence has been assembled as such (Matthews et al. 2018). However, several studies have reported highly-reduced recombination and differentiation that extends along a ∼100 Mb region encompassing the sex-determining M-locus and the centromere. This differentiation was detected as a difference in the read depth of coverage (Matthews et al. 2018) and the frequency of SNPs between males and females (Fontaine et al. 2017). Additionally, several studies have reported sex-specific lethality when genes in the vicinity of the M-locus were recombined or silenced *via* RNAi (Krzywinska et al. 2016; Wood 1990; Mysore et al. 2021b). Moreover, an unexpected mechanism of inheritance bias in males was recently detected for a gene closely-linked to the M-locus that was targeted by a gene drive element (Verkuijl et al. 2022). All these phenomena indicate the sequence divergence of the nominally homomorphic sex chromosome in *Ae. aegypti* warrants deeper characterization. Identification of additional sex-linked sequences outside of a small M-locus would significantly expand the opportunities for further development of genetic sexing and control systems in this major disease vector.

## Results

### Nanopore-based *de novo* genome assembly and its sex-biased diploidization

The first step in our approach is to produce the best initial nanopore-based assembly from a single XY individual. We generated long-read MinION data from an adult male *Aedes aegypti* (specimen T02, Figure 1) and two of its Y-bearing spermatozoa (Figure 1, also see Materials and methods). Given the small size of this organism, only one microgram of native (non-amplified) DNA was available for the MinION sequencing, which is the recommended input amount of DNA for one MinION library that usually yields around 8 Gb of data. Given the genome size of this species, such an output would provide a very low depth of coverage (∼3X) per haploid complement in the diploid assembly (3X ≈ ∼8 Gb/∼1.28 Gb genome size/2 haploid complements). We instead used four times less input DNA (∼250 ng) and sequenced four libraries across four MinION flow cells, yielding a total of 20.1 Gb of data with the read N50 of 16.1 kb. This is expected to achieve ∼8X depth of coverage per haploid complement in the diploid assembly. To test if the addition of reads from the whole-genome-amplified (WGA) DNA would improve the assembly quality (by increasing the depth of coverage), we also generated and sequenced one library with one microgram of male’s WGA DNA, obtaining 6.1 Gb with the read N50 of 8.4 kb. Library from each microdissected Y-bearing (*Nix*-positive) spermatozoon, where the haploid DNA extract underwent necessary WGA, yielded 9.7 and 10.7 Gb, but with a read N50 of only 1.5 kb. Finally, ∼25 ng of the male’s native DNA was used to generate high-accuracy short reads from an Illumina PCR-free library, which gave 86.3 Gb of paired-end data.

**Figure 1.**
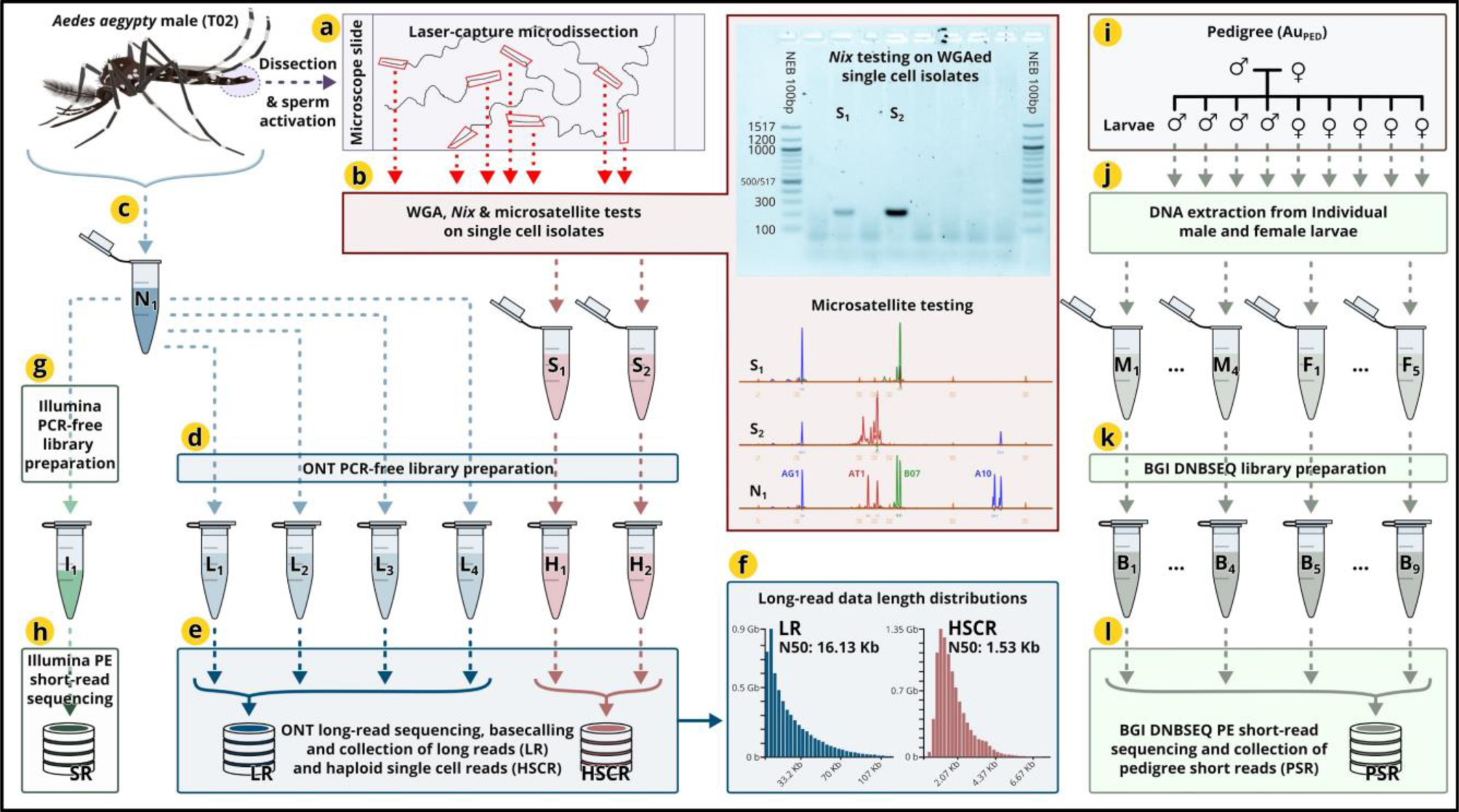
Biological material and sequencing. One *Aedes aegypti* male (T02) was dissected to (a) isolate individual spermatozoa *via* laser-capture microdissection. (b) Spermatozoa isolates then underwent whole genome amplification (WGA), tests for the presence of the partial *Nix* amplicon, and microsatellite genotyping. (c) Total DNA was also extracted from the male’s whole body (minus the dissected *sacculus ejaculatorius*) and used natively (without WGA) to (d) prepare four ONT libraries and (g) one Illumina PCR-free library. DNA from each individual Y-bearing sperm cell isolate (*Nix-*positive S1 and S2, that also showed haploid microsatellite profile) was used to prepare (d) an ONT library. Each ONT library was (e) sequenced on a MinION flow cell, (f) producing long-read data with N50 of ∼16 Kb for the male’s native DNA, and N50 of ∼1.5 Kb for a WGAed single sperm-cell isolate. (i-l) Short-read data from DNBSEQ libraries were generated for four male and five female siblings (single-pair mating, AUSPED; from the same colony as the male T02). The image of a male mosquito is modified from Ramírez, Ana L. (2019): (male) Aedes aegypti. *figshare*. Figure. https://doi.org/10.6084/m9.figshare.7699778.v1. CC BY 4.0.

We ran a series of bioinformatics experiments to produce the most complete *de novo* genome assembly with accurately reconstructed divergent regions. The goal was to achieve the minimal number of missing and the highest number of complete AND duplicated BUSCOs (Diptera_odb10), while having high contiguity statistics (Figure 2). Filipović and colleagues (Filipović et al. 2022) previously demonstrated that the MinION-based assemblies (with low depth of coverage in particular) require polishing with high-accuracy Illumina reads to recover the highest BUSCO scores. Given that the Illumina reads came from the same individual, the polishing removed systemic homopolymer indel errors as well as SNP errors in the initial MinION assemblies. We tested different combinations of sequencing data and assembly parameters and produced ∼70 different initial assemblies. The best initial assembly (T02-5k-v2) was generated with a long-read overlap of 5000 bp and data from the native DNA libraries (long reads for *de novo* assembly and short reads for the assembly polishing). This assembly had a size of ∼1.856 Gb, ∼96.01% complete and ∼46.03% duplicated BUSCOs (fragmented ∼2.13%, missing ∼1.86% BUSCOs), the N50 statistic of ∼245.8 kb, and the L50 of 2070. Interestingly, addition of long-read data from the male’s WGAed DNA did not substantially improve BUSCO results despite increasing the depth of coverage.

**Figure 2.**
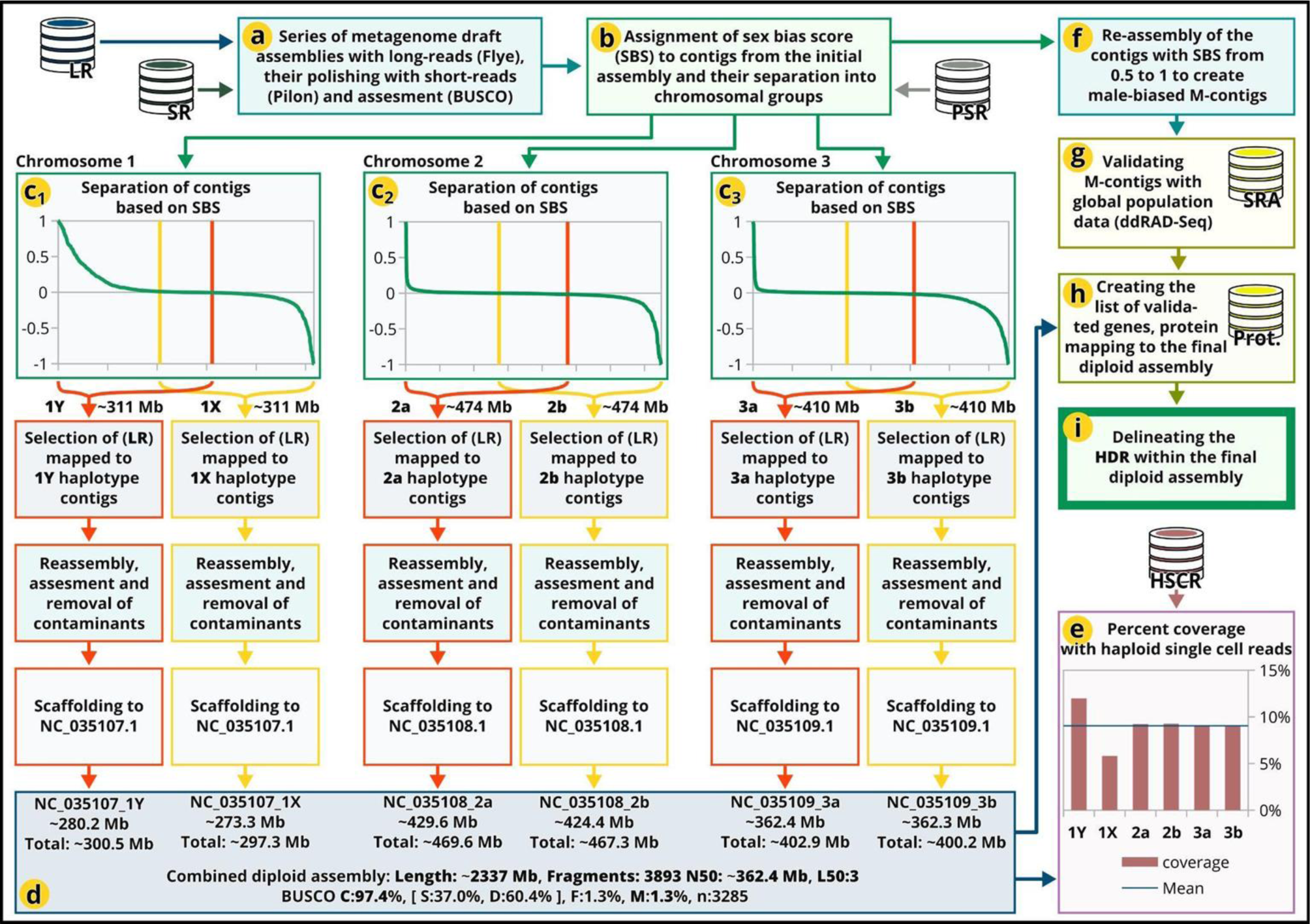
Analytical workflow to identify HDR. The central and left part of the diagram follows a series of steps to obtain the final diploid assembly with the maximized sex-specific differences between the haploid complements. (a) From the series of metagenome assemblies, an initial assembly reflecting the best reconstruction of divergent regions is selected. (b) The assignment of Sex-Bias Score (SBS) enables (c1-c3) the sorting of contigs and their separation into six haplogroups (1Y, 1X, 2a, 2b, 3a, 3b). They are then separately reassembled and scaffolded to produce (d) the final diploid assembly. The right side of the diagram shows the steps to (f) reassemble and (g-h) validate contigs with the highest male bias (M-contigs) and (i) use their genes to delineate HDR. The bottom right corner (e) shows the percent coverage with long-read data from the Y-bearing spermatozoa, where 1Y has twice the percent coverage compared to 1X, while autosomes (2a,b; 3a,b) do not display the percent coverage bias.

To estimate the haploid size (HS) of the best initial assembly (T02-5k-v2), we derived the following formula that considers the percentage of duplicated BUSCOs and assumes that BUSCOs are uniformly dispersed throughout genome:

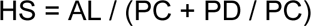

where AL represents the total Assembly Length, PC is the Percent of Complete BUSCOs and PD is the Percent of Duplicate BUSCOs. This gave an estimated genome size of 1.289 Gb. When applied to the BUSCO results for *Ae. aegypti*’s current reference genome assembly (AaegL5.0 (2017d)), the formula produced an estimated size that is very close to the reported size (∼1.276 Gb estimated vs. ∼1.279 Gb reported).

The second step in our approach is to assign the Sex Bias Score (SBS) to each contig in the best initial assembly, which requires the mapping of short reads from at least one male and one female. We define SBS as:

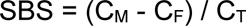

where CM (or CF) is Male (or Female) Coverage that represents a number (or percent) of nucleotide positions on a contig that have a depth of ≥1 of the mapped reads from male(s) (or female(s)). CT is the Total Coverage (or percent) (where male(s) and female(s) mapped short-reads are combined). These statistics should not be confused with the depth of coverage statistics.

In species with male heterogamety (XY males), male-biased (Y-linked) contigs will have SBS value greater than zero (with Y-specific contigs having SBS of 1). Ideally, SBS should be zero for X-linked and autosomal contigs, as percent coverage in males and females is expected to be the same (CM = CF gives SBS of 0). However, depending on the nature of a genome (e.g. the proportion of repetitive regions, haplotype divergence), the nature of available data (heterozygosity and relatedness levels of sequenced male(s) and female(s)), mapping strategy (from strict selection of short reads with the highest mapping score to wider selection with broader range of mapping scores), and depth of coverage, the calculated SBS can take positive and negative values. High SBS values, both positive and negative, reveal contigs with region(s) of high heterozygosity / sequence divergence, and subsequent analyses are needed to pinpoint the sequences of interest.

We calculated SBS for each contig in our best initial assembly (T02-5k-v2) using the whole-genome sequences from four male and five female siblings (pedigree in Figure 1). Their parents (AuPed) originated from the same population as the male (T02) that we used to generate the initial assembly. To note, SBS calculation can be done with the whole-genome sequences from just one male and one female (see further below). Contigs in our initial assembly were also assigned to a given chromosome (1, 2, 3) based on the highest mapping score when aligned to the reference assembly (AaegL5, GCF_002204515.2) (Matthews et al. 2018; 2017d). This produced three groups of chromosome-associated contigs that were then sorted based on their descending SBS value (Figure 2c_1_-2c_3_).

We then generated six haplogroups (1Y, 1X, 2a, 2b, 3a, 3b; Figure 2c_1_-2c_3_) in the following way: contigs associated with chromosome 1 and having a male bias (SBS between 0 and 1) were selected as the Y-associated contigs. To match the chromosome’s size reported in the reference assembly AaegL5, a few additional contigs (SBS slightly below 0) were also added to this group. The X-associated contigs were selected in the analogous way (SBS between -1 and 0, with the addition of contigs with the SBS slightly above 0 to match the reference chromosome 1 size). Haplotype separation was also done for all contigs that were associated with chromosomes 2 and 3.

To further improve the contiguity for each of the six haplogroups, we repeated the assembly process using the original long read data that constituted the selected contigs (Figure 2c_1_-2c_3_). After the BUSCO quality assessment and removal of all potential contaminant sequences (using FCSadaptor (2022) and FCSgx (2022)), each of the six haplogroups (1Y, 1X, 2a, 2b, 3a, 3b) were scaffolded onto a corresponding AaegL5 reference chromosome using RagTag (Alonge et al. 2022). The final product was a diploidized chromosome-level assembly (T02-RagTag-1v3) that maximizes the separation of sex-biased scaffolds. Its high BUSCO statistics (97.4% complete and only 1.3% missing BUSCOs, Figure 2d) were comparable to the latest reference assembly for this species.

We used long-read single-cell data from the male’s Y-bearing spermatozoa (haploid single cell reads, HSCR, Figure 2e) to test if our T02-RagTag-1v3 assembly is correctly phased and scaffolded. If Y and X contain divergent regions and their sequences are correctly separated in an assembly, the reads from Y-bearing haploid spermatozoa are expected to preferentially map to the 1Y scaffold. Indeed, we recovered two times greater percent coverage of the 1Y scaffold compared to the 1X scaffold (Figure 2e). Percent coverage was nearly identical for scaffolds 2a, 2b, 3a and 3b, as is expected for autosomal sequences (Figure 2e).

### Delineation of a region of high divergence between X and Y

We next created an updated version of the best initial assembly (T02-5k-v2) where all contigs with the SBS value greater than 0.5 were re-assembled and marked as male-biased M contigs (T02-M-5k-v2; Figure 2f). This increased the contiguity of male-biased contigs in the updated initial assembly by ∼3 times, with their total sum length of ∼34.08 Mb. To test whether these contigs are consistently detected as male-biased across multiple globally-invasive populations of *Ae. aegypti*, we calculated SBS values using the previously published double-digest RAD sequences (ddRADseq) from males and females originating from Brazil (Rašić et al. 2014b), Indonesia (Rašić et al. 2014b), Vietnam (Rašić et al. 2014b) and Australia (Rašić et al. 2016). To ensure an unbiased mapping of short reads, the entire updated initial assembly (T02-M-5k-v2) was provided for the mapping process. We only selected contigs that were male-biased in all four populations, and this process confirmed 134 contigs (out of 245 initially identified) as male-biased, with their total length summing to ∼24.7 Mb (∼72% of the initially identified length).

We then generated a list of genes found within 134 male-biased M contigs by aligning all protein sequences translated from the reference assembly (AaegL5, GCF_002204515.2) (Matthews et al. 2018; 2017d) using Exonerate (Slater and Birney 2005). This list included the known male-specific genes, *Nix (Hall et al. 2015)* and *myo-sex* (Hall et al. 2014), which supported the accuracy of our workflow. Finally, we mapped the protein sequences of these and all other protein sequences from the *Ae. aegypti*’s annotation (Matthews et al. 2018) to our final diploidized assembly (T02-RagTag-1v3, Figure 2h). This mapping process revealed that nearly all protein sequences from our validated male-biased contigs clustered in the central region of 1Y scaffold, and their gametologs clustered in the central region of 1X scaffold (Figure 3a). Boundaries of this highly divergent region (HDR) on our 1Y and 1X scaffolds were delineated using the start and end positions of the corresponding contigs. Finally, we wanted to test if the same HDR region would be recovered if we used whole-genome sequences from only one male and one female to calculate the SBS in the initial assembly. Indeed, 100% of HDR contigs were confirmed with data from a male and a female originating from a population in Thailand (Rose et al. 2023).

**Figure 3.**
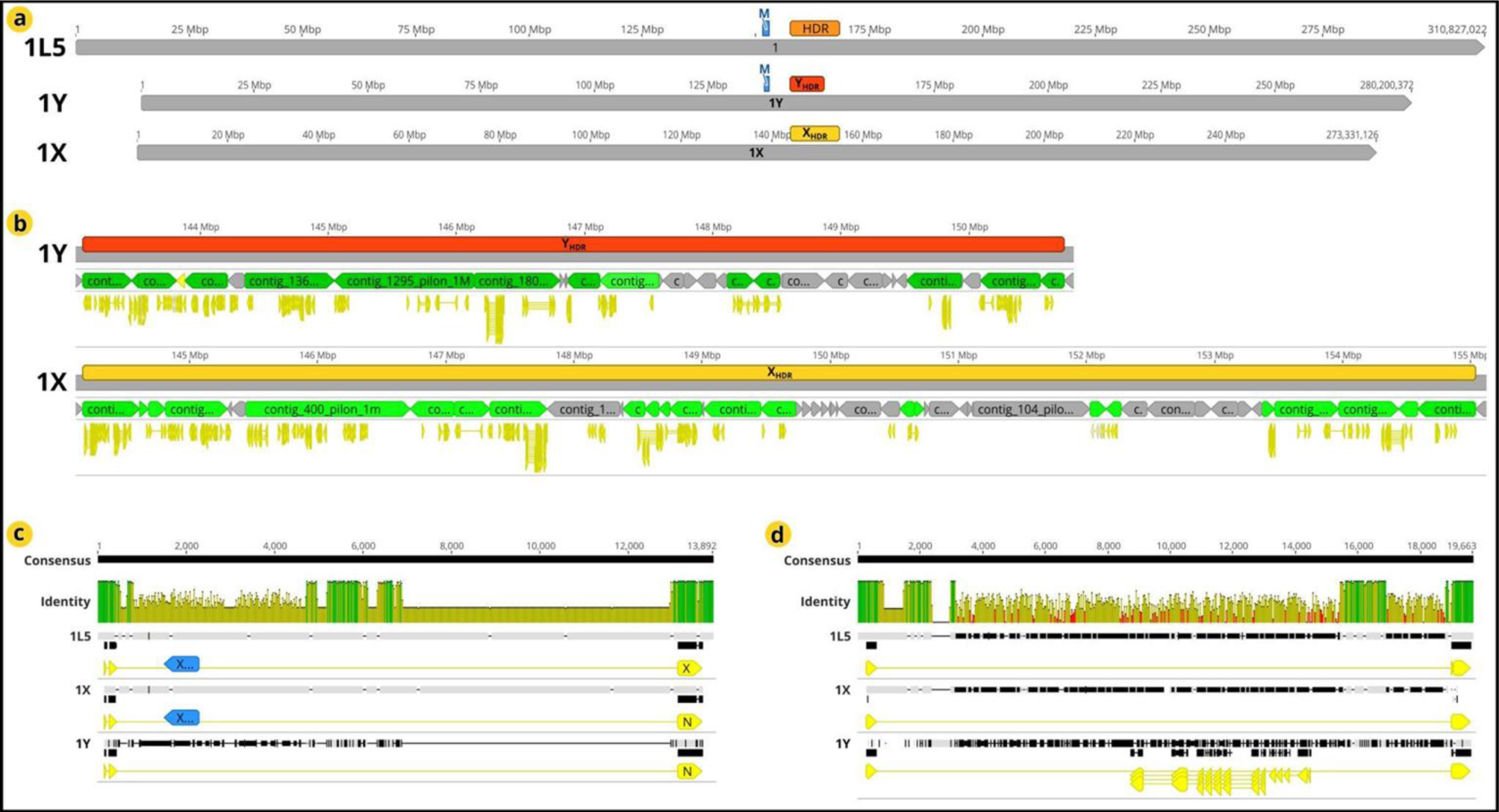
HDR location and structural differences between HDR gametologs. (a) In the reference assembly (AaegL5, (Matthews et al. 2018)), HDR (orange) corresponds to the central part of 1L5 reference scaffold (chromosome 1, NC_035107.1 (2017a)) and is located in the vicinity of the M-locus (light blue). In our diploid assembly, HDR is smaller on the 1Y scaffold (red), and nearly identical in size between the 1X and the reference scaffolds (yellow in 1X, orange in 1L5). (b) An expanded view of the HDR region on 1Y and 1X scaffolds, respectively, showing constitutive contigs (green) and protein CDSs (yellow). (c) An example of an intragenic lncRNA gene (blue) that is found in the intronic region of a protein coding gene in 1X (and in the reference assembly, 1L5), but is absent from the 1Y gametolog. (d) An example of a nested gene CDSs (yellow) found in 1Y but not in the 1X gametolog (or in the reference 1L5).

HDR is situated between 143 Mb and 150.7 Mb in our 1Y scaffold, and between 144.2 Mb and 155 Mb in our 1X scaffold (Figure 3a). In the current reference genome assembly (AaegL5) (Matthews et al. 2018; 2017d) this corresponds to the region between 157.6 and 168.6 Mb on chromosome 1, which is in the centromeric region and ∼5 Mb downstream of the male-determining M-locus (with *Nix* and *myo-sex* genes, Figure 3a). A smaller size of HDR on the 1Y scaffold (∼7.7 Mb) compared to the size on 1X (∼10.9 Mb) is driven by shorter intergenic and intronic sequences (described next). For clarity, we further refer to the HDR sequences and gametologs on 1Y as YHDR, and on 1X as XHDR.

### Divergence patterns between X/Y gametologs in HDR

We found 75 protein-coding genes within HDR, and recovered both gametologs for 58 of them (17 were missing an allele on XHDR). Where present, introns were significantly shorter in the YHDR gametolog (Wilcoxon Signed-Rank Test *W* = 837.5, *p*-value = 0.0012). The median sequence identity between longer introns (>1800 bp) was 41%. For exons, the median number of synonymous substitutions per site (*dS*) between YHDR and XHDR gametologs was 0.027 (maximum 0.096); while the median number of nonsynonymous substitutions per site (*dN*) was 0.003. For comparison, a subset of 72 autosomal single copy orthologs equally spread across autosomal scaffolds (2a, 2b, 3a and 3b) had a median *dS* equal to 0 (maximum 0.194) and a median *dN* equal to 0 (maximum 0.013).

Divergence within HDR is also evident as a differential presence of the long noncoding RNA (lncRNA) genes: out of 13 detected, 12 were found on XHDR (10 intergenic, two intragenic, Figure 3b (left)), and only two on YHDR (one is YHDR-specific, one is shared with XHDR). Another example of divergence in this region is the presence of a nested gene within the YHDR gametolog that was absent from the XHDR gametolog (LOC5568277, Figure 3d).

Sequence divergence in HDR was also apparent as a significantly higher region-wide heterozygosity in males (Mann Whitney U test *W* = 54, *p* = 1.59e-08). The signal of sex-linked differentiation at the HDR loci dominates over the signal of population origin in the analysis of population genetic structure. Namely, individuals from four invasive *Ae. aegypti* populations (Brazil, Vietnam, Indonesia, Australia) were grouped by their sex when HDR loci were used in the discriminant analysis of principal components (i.e. males and females grouped separately, regardless of their population of origin, Figure 4a). Conversely, when loci from chromosomes 2 and 3 were used in the analysis, individuals were grouped by their population of origin, regardless of their sex (i.e. males and females from the same population were grouped together, and all four populations were clearly separated from each other, Figure 4b). Noteworthy are three individuals (two from Vietnam, one from Brazil) that were identified as males by the presence of sequences from *Nix* (the M-locus gene), but had female-specific genotypes at HDR loci (Figure 4a). This pattern indicates recombination between the M-locus and HDR. However, as all three individuals were sampled as larvae, we hypothesize that such recombination may be lethal or incurs high fitness cost at later life stages.

**Figure 4.**
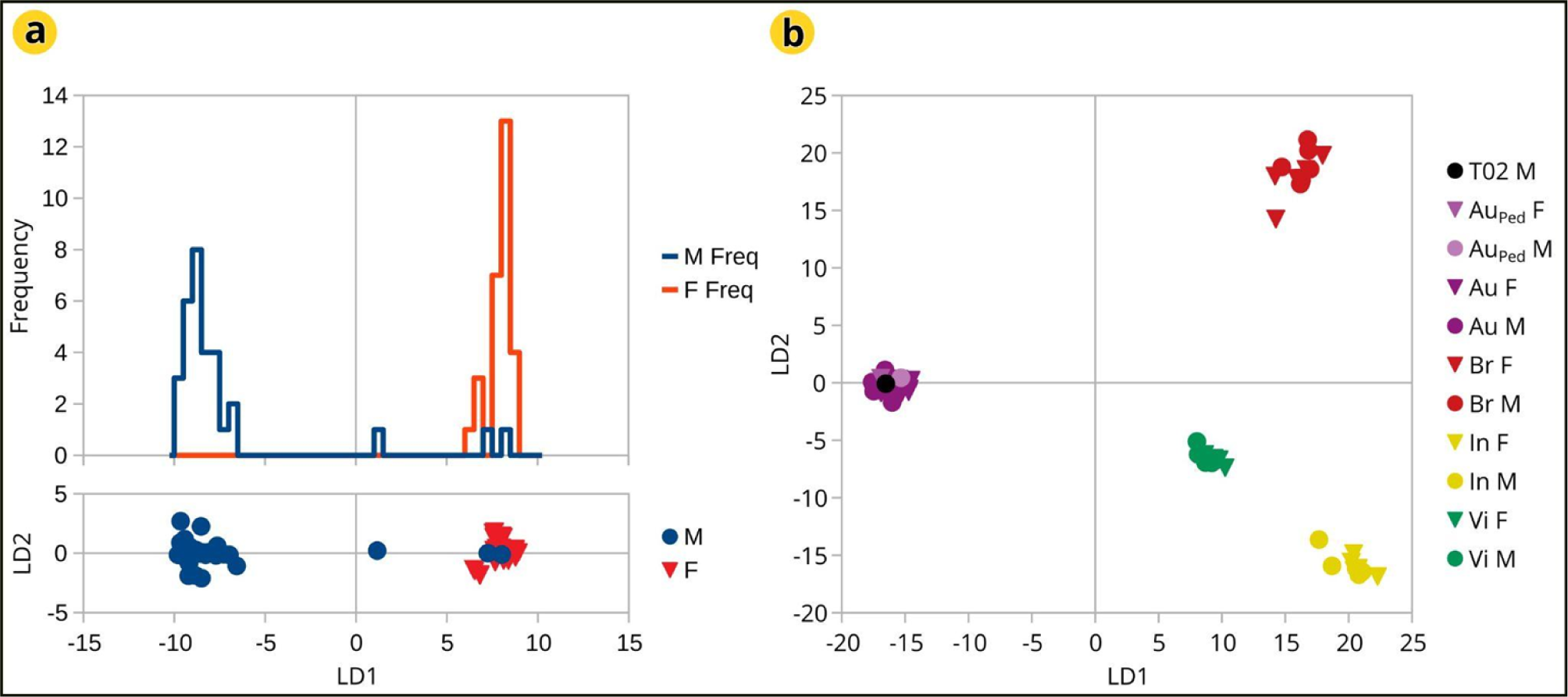
Genetic structuring by sex and geography with HDR and autosomal SNPs. The analysis of genetic structuring shows the X-Y nature of HDR compared to autosomes. (a) When genotypes from HDR are used for the structure analysis (*find.cluster* and DAPC, males (blue) and females (red) are separated into two groups regardless of their population of origin. Noteworthy exceptions are two males that group with females, and one that is located between the two groups. (b) The analysis with autosomal genotypes assigns individuals of both sexes to their population of origin, regardless of their sex (Australia: T02, AuPed, Au; Brazil: Br; Indonesia: In; Vietnam: Vi).

### Recombination within HDR ceased relatively recently

The cessation of recombination between gametologs on sex chromosomes at different time points should generate different gene tree topologies that can be treated as hypotheses for which relative support, such as the expected likelihood weight (*ELW*), can be calculated (Zhang et al. 2022). We used the *ELW* approach to test four hypotheses (tree topologies, blue and red in Figure 5a). The primary hypothesis (blue) is that the cessation of recombination between YHDR and XHDR is very recent, occurring after the split between populations (i.e. split between XHDR in the Australian and XHDR in the reference LVP strain that originated from West Africa). Secondary group of hypotheses (red) are that the cessation of recombination between YHDR and XHDR predates the population split; three alternative scenarios (different shades of red, Figure 5c) consider whether XHDR/YHDR split occurred before, during or after *Ae. aegypti* diverged from its nearest described relative, *Ae. mascarensis*. The analyses were done separately for each HDR gene where an ortholog from a distantly-related *Ae. albopictus* could be identified (48 out of 75 HDR genes). Density plot for the *ELW* values shows that the group of secondary hypotheses (split between YHDR and XHDR predates the population split) is strongly supported (average *ELW* > 0.93) for the majority of tested genes (red in Figure 5a).

**Figure 5.**
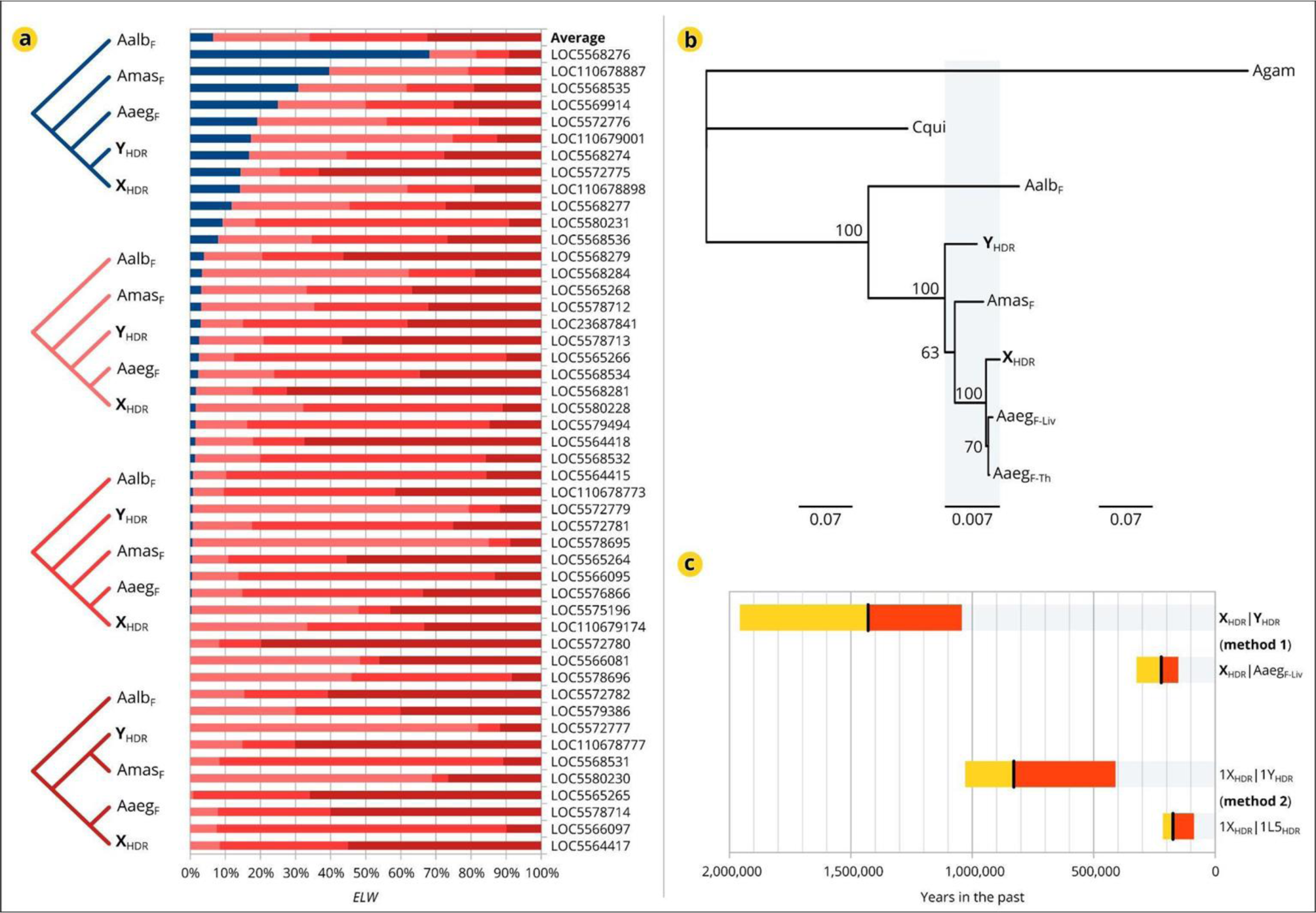
Timing of the split between HDR gametologs. (a) Four alternative hypotheses (tree topologies) were tested: (blue) YHDR and XHDR diverged very recently *versus* YHDR and XHDR diverged after (light-red) or before (mid-red) or during (dark-red) the split between *Ae. aegypti* (AaegF) and *Ae. mascarensis* (AmasF). The hypothesis of a very recent XHDR/YHDR divergence (blue) is well supported (*ELW* >50%) for only one gene. (b) ML tree for the concatenated exonic sequences of 48 genes from *Ae, aegypti*’s HDR and their orthologs in *Ae. albopictus* (Aalb)*, Cx. quinquefasciatus* (Cqui)*, An. gambiae* (Agam). The tree is “blown up” 10 times in the light-blue section for better visibility. (c) Dated cessation of recombination in HDR (years in the past).

To date the recombination cessation in HDR, we used two approaches. The RealTime-Branch Length analysis in TimeTree (method 1), that used the tree topology inferred with IQ-Tree from the concatenated exonic sequences (Trifinopoulos et al. 2016) (Figure 5b), gave the estimate between 1.04 and 1.96 million years since YHDR and XHDR last recombined (method 1, Figure 5c). This range overlapped with the estimate of the split time between *Ae. aegypti* and *Ae. mascarensis* (0.81-1.50 MYA). In contrast, recombination cessation between XHDR gametologs from different *Ae. aegypti* populations (Australia and the Liverpool strain) was estimated as much more recent: 0.15-0.32 MYA. In the approach where both coding and non-coding sequences were used (method 2, Figure 5c), the recombination cessation between YHDR and XHDR was estimated at 0.41 and 1.03 MYA, while the recombination cessation between XHDR from different populations was estimated at 0.09 and 0.22 MYA. Importantly, the split time between X/Y gametologs in HDR did not overlap with the split time between XHDR alleles across populations for any combination of parameter values (the number of generations per year, mutation rate, split time between *Ae. aegypti* and *Ae. albopictus*).

### Gene expression in HDR is female-biased in adults

Analysis of sex-biased gene expression across different life stages (larvae, pupae and adults) revealed that HDR genes have significantly female-biased expression in adults, but not in larvae and pupae (Figure 6). The distribution of values for the normalized ratio of expression (transcripts per million, TPM) in females and males (log2(TPMF / TPMM)) was compared between genes in HDR, autosomes (chromosomes 2 and 3, Aut.) and pseudoautosomal region (Par., the first 50 Mb of the sex-determining chromosome 1). In larval and pupal stages, expression profile for the HDR genes was not different from other genome parts (permutation test of equality of densities, larvae: HDR vs. Aut. *p* = 0.558; HDR vs. Par., *p* = 0.656, Figure 6a; pupae: HDR vs. Aut. *p* = 0.179; HDR vs. Par., *p* = 0.164, Figure 6b). A significant shift towards female-biased expression of HDR genes was evident in adults (HDR vs. Aut. *p* = 0.016; HDR vs. Par. *p* = 0.005, Figure 6c).

**Figure 6.**
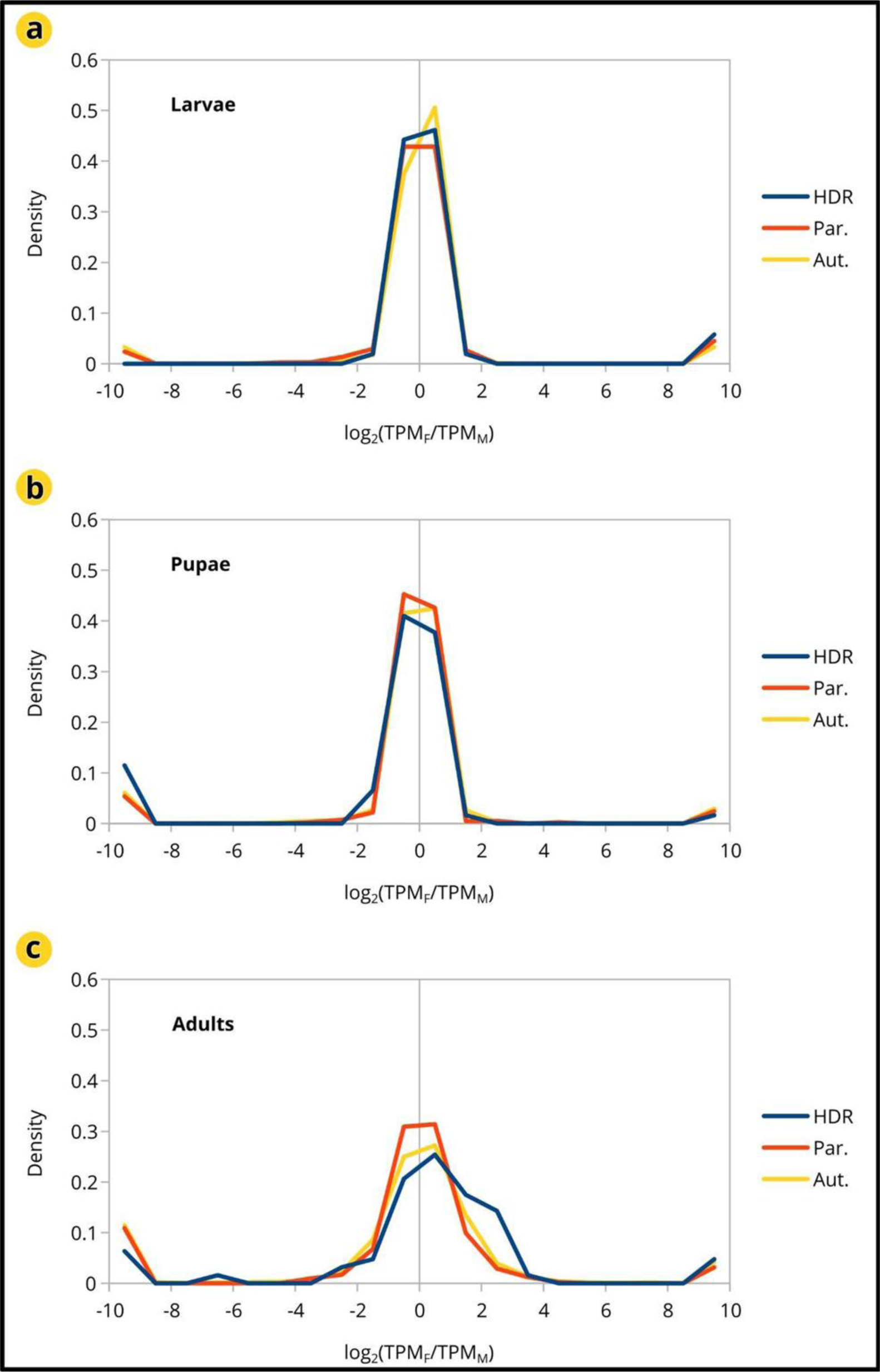
Sex-biased expression across life stages and genomic regions. (a) In larvae and (b) pupae, the distribution of normalized values for the gene expression difference between males and females overlaps for HDR, autosomal (Aut.) and pseudoautosomal (Par.) genes. Gene expression is not sex-biased for either region. (c) In adults, gene expression is significantly female-biased for HDR genes, but not for Aut. and Par. genes.

This female-biased expression is largely driven by genes that are overexpressed in ovaries. Namely, the proportion of ovary-biased genes is significantly higher in HDR (0.267, CI: 0.203-0.361) than in the autosomes (0.125, CI: 0.119-0.131) and the pseudoautosomal region (0.079, CI: 0.054-0.110) (confidence intervals do not overlap between HDR and Aut. Or Par.). Conversely, the proportion of testis-biased genes is not statistically different between HDR (0.147, CI: 0.085-0.243), the autosomal region (0.143, CI: 0.137-0.150) and the pseudoautosomal region (0.106, CI: 0.079-0.139). Therefore, HDR is enriched for ovary-biased genes while not being depleted for testis-biased genes. The same trend was observed for genes that are expressed in the adult brain, with this female bias being statistically significant compared to the autosomes but not to the pseudoautosomal region (HDR 0.067, CI: 0.042-0.134; Aut. 0.034, CI: 0.031-0.038; Par. 0.036, CI: 0.024-0.055).

## Discussion

We demonstrate an approach that can recover haplotypes of homomorphic sex chromosomes in a nanopore-based assembly from a single XY male. Separation of haplotypes is guided by the Sex-Bias Score (SBS) that quantifies differential mapping of short reads from males and females, and the aligning of protein sequences from the male-biased contigs (SBS > 0) onto the diploid assembly enables identification of X/Y gametologs. We applied the approach to *Ae. aegypti*, a mosquito with a large repetitive genome and a homomorphic sex chromosome system (Matthews et al. 2018), where previous studies demonstrated SNP differences between males and females and a reduced recombination on the sex chromosome around the male-determining locus (M-locus) (Matthews et al. 2018).

We focused on the X/Y sequences with very high divergence (SBS > 0.5) and recorded their clustering near the centromeric region of *Ae. aegypti*’s chromosome 1, approximately 5 Mb downstream of the male-determining M-locus. We delineated this highly divergent region (HDR), noting a ∼3 Mb difference in size between homologous chromosomes (∼10.9 Mb on XHDR vs. ∼7.7 Mb on YHDR, Figure 3a-3b). XHDR and YHDR harbour divergent alleles of protein coding genes (median *dS =* 0.027, *dN* = 0,003; shorter introns on YHDR), most lncRNA genes have only X allele, and one nested protein-coding gene (within LOC5568277) is Y-specific (Figure 3c). Until now, all these differences have been invisible, as the reference genome assembly for this species (AegL5 (2017d)) had alternative haplotypes merged for the sex-determining chromosome 1 the same way as for the two autosomes (Matthews et al. 2018).

Accurate separation of haplotypes on homomorphic sex chromosomes helps explain some surprising experimental results in this mosquito. Mysore and colleagues recently reported that the silencing of some lncRNA genes within and adjacent to the male-determining M-locus resulted in significant female larval mortality, often without an apparent fitness cost to males (Mysore et al. 2021a). These results are unexpected, as female-specific mortality should not be observed when Y-specific (M-locus) genes are silenced. The authors (Mysore et al. 2021a) noted that such results could be due to off-site targeting of genes that are missing from the *Ae. aegypti*’s reference assembly, possibly within a known gap in the reconstructed M-locus sequence. In contrast to the reference assembly where some of the targeted genes have been annotated as Y-specific (Matthews et al. 2018), our assembly contains their sequence outside of the M-locus and alleles on Y and X scaffolds, which permits the possibility of allele-specific (and sex-specific) effects of their silencing. Our assembly also reveals that most lncRNA genes in the newly-evolved HDR have become X-specific, and this “decoupling” from the lncRNA genes in the M-locus could indicate their opposite sex-specific effects.

The M-locus has been considered as the only Y-like genomic region (∼1.5 Mb long) in this species, and as such, represents an attractive target for different genetic constructs (Burt and Deredec 2018; Condon et al. 2007; Gamez et al. 2021). However, it may not be easily accessible to the transgene integration technology. For instance, Navarro Paya D. (Navarro Paya 2017) described unsuccessful attempts to integrate an exogenous DNA sequence into the *Nix* gene, a male-determining factor in the M-locus. We now report ∼10 Mb of putative YHDR- and XHDR-specific targets that should significantly expand the opportunities for engineering the sex-targeting systems to control *Ae. aegypti*.

Split between YHDR and XHDR predates the previously estimated time of *Ae. aegypti’*s movement into continental Africa (<120,000 ago, Bennett et al. 2016). Our estimated divergence time between XHDR in the Australian and the reference strains (that approximates divergence time between ancestral populations) ranges between ∼90,000 and ∼320,000 years (methods 1 and 2, Figure 5c). Divergence time between XHDR and YHDR is greater (0.41-1.96 MYA, estimated across two methods, Figure 5c) and overlaps with our estimate of the split time between *Ae. aegypti* and *Ae. mascarensis* (0.81-1.5 MYA). Soghigian and colleagues reported a much older split time for the two sister species (4-15 MYA) based on the analyses of a few nuclear and mitochondrial genes (Soghigian et al. 2020). As we used 48 HDR genes with the sex-linked evolutionary history, it is possible that the discrepancy in the absolute time scale may be due to different molecular clocks for genes on autosomes and genes on sex chromosomes. Regardless of the absolute time (in years), the overlap between the time of speciation and time of recombination cessation in HDR is noteworthy. Namely, hybridization patterns between *Ae. aegypti* and *Ae. mascarensis* follow the Haldane’s speciation rule for male hybrid sterility (Presgraves and Orr 1998) and have been attributed to genes that are linked to, but not identical with, the M-locus (Hilburn and Rai 1982). Recent theoretical work predicts such hybridization effects for genes in the newly-diverged regions of homomorphic sex chromosomes (Mrnjavac et al. 2023).

In addition to the sequence data, gene expression data further supports the hypothesis of an evolutionarily young sex-linked region: HDR is enriched for ovary-biased genes but is not depleted for testis-biased genes compared to autosomes and the pseudoautosomal region of the sex chromosome. This pattern (enrichment for ovary-biased genes, and retention of testis-biased genes) has been reported in other taxa, including another insect pest, the mountain pine beetle *Dendroctomus ponderosae* (Bracewell et al. 2017), and is interpreted as evidence for the early stages of feminization in the neo-X chromosome (Bracewell et al. 2017).

In conclusion, our approach for identifying divergent sequences of homomorphic sex chromosomes expands the toolbox for studying “cryptic” sex chromosome structures in non-model species. Applied to the deadly mosquito *Ae. aegypti,* it generated original insight into the sequence and gene expression patterns in a newly-diverged region within a homomorphic sex chromosome system. Megabases of previously invisible sex-linked sequences provide new putative targets for engineering the genetic sexing systems and sex ratio distorters to control this invasive vector.

## Materials and methods

### DNA isolation from a single mosquito and its individual sperm cells

A male *Ae. aegypti* from a laboratory colony of Australian origin (Cairns colony, QIMR Berghofer MRI) was separated upon emergence for 3 days and dissected in the insect physiological solution. The dissected male was then placed in 100% ethanol, and the extracted *sacculus ejaculatorius* was transferred onto a sterile microscopy slide with 1000 μL of a sperm activating solution described by Pitts *et al*. (Pitts et al. 2014). The puncturing released and activated the spermatozoa to disperse away from each other. Dispersed spermatozoa were pipetted on to Zeiss microdissection slide where ∼2000 μl of additional activating solution was added and left to air-dry for 24 hours at room temperature (RT). After air-drying for 24 hours at RT, the dispersed spermatozoa were individually microdissected using the laser-capture Zeiss microscope. Each extract underwent whole genome amplification (WGA) using the Qiagen Repli-G Advanced single-cell kit, and then treated with S1 endonuclease (Thermo Scientific) following the manufacturer’s protocol. The amplified single-cell extracts were tested for diploid contamination using a set of five microsatellite loci (as in (Rašić et al. 2014a)), as well for the presence of the *Nix* and *myo-sex* amplicons. Two extracts that showed a clear haploid profile (homozygosity at microsatellite loci that were heterozygotic in the donor male) and the presence of the M-locus (positive for the partial amplicon of the *Nix* gene) were used for the preparation of two ONT libraries (one for each individual Y-bearing spermatozoa). DNA from the dissected donor male was extracted using a customized bead-extraction protocol (as in Filipović *et al*. (Filipović et al. 2022)), obtaining a total of 1.1 μg of high-molecular weight (HMW) DNA.

### Long-read and short-read sequencing

Four ONT libraries (2023a, 2023b, 2023c, 2023d) were prepared with HMW DNA from the single male donor, following the manufacturer’s guidelines for the Ligation Sequencing Kit SQK-LSK109 (Oxford Nanopore Technologies, Cambridge UK), with the exception of using ∼250 ng as the starting material for each library. One additional ONT library was prepared with the recommended amount of input DNA, where ∼20 ng of male’s DNA was whole-genome-amplified using the Qiagen Repli-G kit. Two ONT libraries (2023f, 2023g) were prepared for two individual Y-bearing spermatozoa. A total of seven ONT libraries were sequenced on seven R9.4.1 flow cells using the MinION sequencing device and the ONT MinKNOW Software (Oxford Nanopore Technologies, Cambridge UK).

One Illumina PCR-free library (2023e) was prepared with ∼25 ng of native (non-amplified) DNA from the male donor and sequenced in one NextSeq 550 flow cell using the 300 bp PE chemistry. Nine DNBSeq libraries were prepared with one microgram of DNA from each individual larva (four males and five females) and sequenced using the 200 PE chemistry.

### Genome assembly and the alignment of long- and short-read data

The basecalling for all ONT reads (seven libraries) was done with Guppy v.6.0.1, using the “super accurate mode” (minimum quality 7). All genome assemblies were produced with the long-read data from the male donor using Flye v.2.9 (Kolmogorov et al. 2019), with an increasing overlap length (1 kb-10 kb) in increments of 500 bp (as described in Filipović *et al*. (Filipović et al. 2022)), using the metagenomic mode. The polishing of each assembly was done with Pilon (Walker et al. 2014) using the high-accuracy Illumina data from the male donor (T02). The quality assessment of each assembly was done with BUSCO (Simão et al. 2015) using the fly gene set Diptera_odb10.

The short-read data (DNBSeq and Illumina ddRADseq) from individual males and females and their pooled data were aligned to the assembly versions with Bowtie2 (Langmead and Salzberg 2012) using the --very-sensitive mode. Given that the Sex Bias Score is sensitive to the variability in the number of mapped reads (used to calculate the percent coverage), only data with a very high mapping quality (MQ>=40) were used. Additionally, for the ddRADseq data we considered positions with more than one read per position (depth of >1) when calculating SBS, given the greater noise of read depth with possible allelic dropout for double-digest libraries. For the same reason, we also chose contigs with SBS > 0.05 (rather than SBS > 0) as male-biased when using ddRADseq data. The alignment of long-read data and the assembled contigs was done using minimap2 (Li 2018) with default parameters. The scaffolding of contigs was done with RagTag (Alonge et al. 2022) in the scaffold mode, with -q 0 -f 100 –mm2-params -x asm5. The long-read data from single spermatozoa were aligned to the assemblies using minimap2 (Li 2018) and only data with the highest mapping quality (MQ=60) were used for downstream analyses.

### Identification of gametologs

Protein-coding genes were identified by aligning the translated coding sequences (all isoforms) of each gene from the existing *Ae. aegypti* genome annotation to the male-biased contigs with Exonerate (Slater and Birney 2005) using the parameters: -model protein2genome - bestn 30. Only proteins for which at least 90% of their sequence length was aligned to the heavily male-biased contigs were retained for further analyses. To identify gametologs, these sequences were aligned to the entire diploidized genome where ≥ 50% of the sequence length needed to be mapped. If a coding sequence had multiple alignments on Y- or X-linked contigs, we retained those with the lowest number of synonymous substitutions per synonymous sites (*dS*), in order to limit the analyses to true gametologs rather than to paralogs. Orthologs in two other Culicinae, *Ae. albopictus* and *Culex quinquefasciatus,* as well as the malaria mosquito *Anopheles gambiae* (Anophelinae) were identified from Vectorbase (Giraldo-Calderón et al. 2022) and through a BLASTX search on a local database that contained the translated sequences from the latest genome annotations for *A. gambiae*, *A. albopictus* and *C. quinquefasciatus*, using the *e*-value cutoff of 10^-10^.

### Genetic structure analysis

To test the genetic structuring based on the genotypes at the HDR and autosomal loci, we used the short-read ddRAD-seq data from males and females originating from Brazil (Rašić et al. 2014b), Indonesia (Rašić et al. 2014b), Vietnam (Rašić et al. 2014b) and Australia (Rašić et al. 2016), as well as the whole-genome Illumina data for the male donor (T02) and the whole-genome DNBseq data for the parents from the pedigree sample (AuPed, Figure 1). The total number of analysed individuals was 59 (28 females and 31 males). Data processing, sequence mapping and realignment, and genotype calling were done following the steps outlined by Igor Filipović (Filipović 2023). The final number of SNP loci was 1,921 for HDR, and 4,052 for the autosomes (chromosomes 2 and 3), which had ≤15% missingness per individual. Discriminant analysis of principal components (DAPC) was done on the groups inferred via the *find.cluster* function in the R package adegenet (Jombart 2008) (i.e. the groups were not defined *a priori*).

### *dN/dS* calculations and timing of the recombination cessation in HDR

Clustal Omega v1.2.3 implemented in Geneious (Biomatters development team 2020) was used to obtain pairwise alignments for the coding sequences, and the *dN* and *dS* values between gametologs and orthologs in *Ae. albopictus* and *C. quinquefasciatus* were calculated using the Nei–Gojobori method (Jukes–Cantor) in MEGA 11 (Tamura et al. 2021) with gamma variation among sites.

The relative dating of the recombination cessation between gametologs in HDR was done using the hypothesis testing framework (Zhang et al. 2022), where the expected likelihood weight (*ELW*) was calculated for four tree topologies (four hypotheses). For each gene, the alignment included CDS from the XHDR and YHDR gametologs from the T02 male (Australian population) and from the X-linked orthologs in the *Ae. aegypti* Liverpool strain, *Ae. mascarensis* and *Ae. albopictus.* Each CDS consensus for the X-linked orthologs was formed with the reads from females (XX) only. For the *Ae. aegypti*’s Liverpool strain (2014a, 2014b; Hall et al. 2014) and *Ae. mascarensis* (2020a, 2020b; Rose et al. 2020), the reads were mapped to the AaegL5.0(2017d) reference assembly (denoted as AaegF). For *Ae. albopictus* (2017b, 2017c), the reads mapped to Aalbo_primary.1(2019) reference assembly (denoted as AalbF). The Newick tree format for the hypothesis of a “recent recombination cessation” is (AalbF,(AmasF,(AaegF,(YHDR,XHDR)))); and the hypotheses of “older recombination cessations” are (AalbF,(AmasF,(YHDR,(AaegF,XHDR)))): (AalbF,(YHDR,(AmasF,(AaegF,XHDR)))): and (AalbF,(YHDR,AmasF),(AaegF,XHDR));. The *ELW* values were computed using the approximately unbiased (AU) topology testing in IQ-Tree web server (Trifinopoulos et al. 2016) with 10,000 resamplings.

The molecular dating of the XHDR-YHDR split was done in two ways. Method 1 employs the TimeTree analysis (RealTime-Branch Length) in Mega 11 (Tamura et al. 2021). The analysis was done with the concatenated CDSs from 48 HDR genes where orthologs could be identified in *Ae. albopictus*, *Cx. quinquefasciatus* and/or *An. gambiae.* We also included the concatenated consensus CDSs from *Ae. aegypti* females originating from Thailand (denoted as AaegF-Th). The maximum likelihood phylogeny was generated in the IQ-TREE web server (ref) using the analytically-determined substitution model, FreeRate heterogeneity and ultrafast bootstrap analysis with 10,000 bootstrap alignments. The resulting tree in the Newick format was provided to the RealTime algorithm; *An. gambiae* and *Cx. quinquefasciatus* were used as an outgroup; the calibration constraint for the split time between *Ae. albopictus* and *Ae. aegypti* was based on the confidence interval (25-38 MYA) reported by Soghigian and colleagues (Soghigian et al. 2023). For the method 2 (*t = D/2μ)*), genetic distance *D* was calculated for the entire aligned region (coding and non-coding + intergenic sequences) in Geneious (Biomatters development team 2020), and the range of value for mutation rate *μ* (3.0-1178.5 x 10^-9^) and the number of generations per year (10-15) were based on the previously-reported estimates (Rose et al. 2023).

### Differential gene expression analysis

Differential expression of genes in HDR was characterized using the published datasets that contain quantified RNA-seq reads (as Transcripts per million, TPM) for the replicate libraries across different life stages and tissues (larva, early and late pupa, male and female adult carcass, ovary, testis, brain (Supplemental Data 7 and 8 from Matthews *et al*.(Matthews et al. 2018)). A gene was considered ovary-biased (or testis-biased) if TPM in the ovary (or testis) dataset was at least two times greater than TPM from the female (or male) carcass and the first instar larva (L1) dataset. Early developmental stages such as L1 have been considered a suitable reference for somatic non-sexual tissues when assessing gonad-specific expression in *Ae. aegypti* (Whittle and Extavour 2017). Sex-biased expression was analyzed using the logarithmized TPM ratio between female and male dataset (log2(TPMF / TPMM)) from the sexed larvae and pupae, as well as adult gonads, carcass and brain for each sex. Statistical significance of difference between densities for the log2(TPMF / TPMM) data was determined with a bootstrap hypothesis test of equality (*sm.density.compare*) in the R package *sm* (Bowman and Azzalini 2021), using 1,000 permutations.

### Data Access

The sequencing data generated in this study have been deposited to the NCBI and are openly accessible as SRA experiments SRX20723278-SRX20723284 under the BioProject accession number PRJNA946909.

## Acknowledgments

IF, GR and JMM were supported by the NIH R01 AI143698-01A1 grant awarded to JMM and GR. IF was also supported by The University of Queensland Graduate School Research Training Program Tuition Fee Offset and Research Training Program Stipend scholarship. The work was supported by the QIMR Berghofer MRI Seed Grant and BGI PRP awarded to GR, the NIH R01 AI143698-01A1 grant awarded to JMM and GR, and the core funds of the Mosquito Control Laboratory at QIMR Berghofer MRI. The authors would like to thank Paul Collins, Nigel Waterhouse and Macky Edmunson from QIMR Berghofer MRI for technical assistance.

## Authors contribution

IF and GR designed the study, GR and JMM provided resources, IF performed the experiments, IF and GR analyzed the data, IF and GR wrote the draft manuscript, GR and JMM edited the manuscript.

